# Dual role of Bnl/Fgf signaling in proliferation and endoreplication of Drosophila tracheal adult progenitor cells

**DOI:** 10.1101/393306

**Authors:** Cristina de Miguel, Josefa Cruz, David Martín, Xavier Franch-Marro

## Abstract

**Abstract:** Adult progenitor cells activation is a key event in the formation of adult organs during development. The initiation of proliferation of these progenitor cells requires specific temporal signals, mostly of them still unknown. In *Drosophila*, formation of adult tracheal system depends on the activation of tracheal adult progenitors (tracheoblasts) of Tr4 and Tr5 tracheal metamers specific spiracular branches (SB) during the last larval stage. The mitotic activity of these tracheoblasts generate a pool of tracheal differentiated cells that migrate during pupal development along the larval trachea by the activation of the Branchless (Bnl)/Fibroblast growth factor (FGF) signaling to form the abdominal adult tracheal system. In here, we found that, in addition to migration, Bnl/FGF signaling, mediated by the transcription factor Pointed, is also required for adult progenitor cell proliferation in the SBs. Moreover, we found that tracheoblast proliferation in Tr4 and Tr5 SBs relies on the specific expression of the FGF ligand *Bnl* in their nearby transverse connective branches. Finally, we also show that, in absence of the transcription factor Cut (Ct), Bnl/FGF signaling induces endoreplication of differentiated tracheoblast daughter cells by in part promoting *Fizzy-related* (*Fzr*) expression. Altogether, our results suggest a dual role of Bnl/FGF signaling in tracheal adult progenitors, inducing both proliferation and endoreplication of tracheoblasts in late larval development, depending on the presence or absence of the transcription factor *ct*, respectively.

**Author summary:** The generation of adult organs and tissue renewal are complex processes that depend on the proliferation and posterior differentiation of undifferentiated progenitor cells in a temporal coordinated manner. Although many signals that regulate the activity of progenitor cells have been identified, the characterization of the mechanisms underlying the temporal and spatial control of such events remain unknown. The tracheal system of *Drosophila*, the respiratory organ, forms during embryogenesis and it is remodeled during metamorphosis from quiescent adult progenitor cells that proliferate. We have discovered that this proliferation depends on the activation of the FGF signaling as mutations that either inactivate or over-activate the pathway blocks cell division or induced over-proliferation of progenitor cells, respectively. Interestingly, we have found that the same signaling pathway also controls tracheal progenitor cells differentiation by promoting endoreplication. We found that this dual role of FGF signaling in adult progenitor cells, depends on the presence or absence of the transcription factor Cut. Altogether, our results, reveal the mechanism that control the division and differentiation of progenitor cells and open the possibility that analogous signaling pathway may play a similar role in vertebrate stem cell regulation and tumor growth.

## Introduction

The formation of adult organs depends on the activation of progenitor undifferentiated cells during development. Temporal regulation and level of activity of adult progenitor cells are critical to coordinate their proliferation and differentiation in order to form an adult functional tissue. Although great progress has been achieved in the identification of signals that regulate the activity of progenitor cells, the characterization of the mechanisms underlying the temporal and spatial control of such events remain far from understood. Here, we use the formation of the adult tracheal system of *Drosophila*, the tubular organ responsible for oxygen transport [1,2], to address this issue.

The embryonic trachea of *Drosophila* develops from 10 bilaterally symmetric clusters (Tr1-Tr10) of ~80 cells that invaginate to form epithelial sacs that remain connected to the epidermis through the spiracular branches (SBs) [3]. These cells migrate and differentiate under the control of the Fibroblast growth factor (FGF)/branchless (Bnl) signaling pathway during embryogenesis to generate a network of interconnected tubes that will function as the larval tracheal system. This larval tracheal network is then heavily remodeled during pupal metamorphosis from a reduced number of different adult precursors cells, called tracheoblasts [2,4-14]. One type of these cells are the abdominal SB tracheoblasts, which are multipotent undifferentiated cells that are specified in the embryo and remain quiescent until the third larval instar (L3), when they proliferate and differentiate to form the definitive adult abdominal tracheal system [2,5,6,13-15]. Remarkably, although SB tracheoblasts are present in all abdominal metameres of the larvae (Tr4-Tr9), only those from the Tr4 and Tr5 metameres proliferate and differentiate during metamorphosis to generate the definitive adult abdominal airways [13,15]. However, the molecular mechanisms underlying such spatially-restricted SB tracheoblast proliferation remains elusive.

Upon activation, tracheoblasts in the Tr4 and Tr5 SBs start to proliferate. However, tracheoblast mitotic activity does not occur uniformly. Instead, four cell populations with different proliferation rates can be distinguish in the SB [13,15]. The tracheoblasts located in the intermediated SB zone, called zone 2, presents the higher rate of proliferation. After division, these cells move towards zone 1, the most dorsal part of the SB closest to the DT, and instead of mitosis, initiate one round of DNA replication by activation of the anaphase promoting complex/cyclosome (APC/C) activator Fizzy related (Fzr) to become 4C at the wandering stage [13]. Finally, tracheoblasts at zone 3 presents a very low mitotic rate, while those located at zone 4, at the most ventral tip of the SB, do not proliferate [13]. Previous work has shown that the difference in the proliferation rate of the SB tracheoblasts depends on the relative abundance of the homeobox transcription factor Cut (Ct). Thus, whereas the highly proliferative zone 2 requires intermediate Ct amounts, the non-proliferative zone 1 demands the complete absence of Ct [15]. This work also shows that the different levels of Ct result from the positive and negative regulatory activity of Wingless and Notch signaling pathways, respectively [15]. This particular expression pattern of Ct, however, is not only detected in the Tr4 and Tr5 SBs, but in all abdominal SBs, from Tr4 to Tr9, thus suggesting that other factors must act for the spatial control of the proliferation and differentiation of abdominal SB tracheoblasts.

To address this question, we focus our attention to the Fgf/Bnl signaling pathway, as it has been shown that the FGF receptor Breathless (Btl) is expressed in the endoreplicative cells of zone 1 and in the proliferative growth zone 2 of all SBs [13]. This expression of Btl has been linked to the migration of the Tr4 and Tr5 tracheal progenitors into the posterior part of the abdomen later on pupal development [16]. Here, we found that Fgf/Bnl signaling also exerts a dual regulatory role in the control of tracheoblast development. First, we showed that activation of Bnl/Fgf signaling in the Tr4 and Tr5 SBs is required to initiate and promote tracheoblast proliferation at zone 2. Remarkably, we show that the spatially restricted tracheoblast proliferation to Tr4 and Tr5 SBs is due to specific expression of the Fgf ligand Bnl in those metameres. In addition, we showed that Fgf/Bnl signaling also promotes endoreplication in differentiated SB progenitor cells that express Fzr at zone 1. Finally, we demonstrate that the dual regulatory effect induced by Bnl/Fgf is transduced via the transcription factor pointed (Pnt). Altogether, our results demonstrate that the Fgf/Bnl pathway is critical in SB tracheoblasts development, playing a dual role on promoting mitotic cell division as well as cell growth through endoreplication.

## Results

### Bnl/Fgf signaling promotes cell proliferation in SB cells

In order to analyze the role of Bnl/Fgf signaling in SB development we either overactivate or inactivate the pathway in adult progenitor cells at L3. Depletion of the FGF ligand Bnl in all tracheal cells using the *btlGAL4* driver completely abolished proliferation of SB progenitor cells (Fig. 1A-B). Consistently, overexpression of Bnl using the same driver promoted over-proliferation of SB progenitor cells and the overgrowth of the SB (Fig. 1 A and D). Interestingly, we found that overexpression of Bnl induces overgrowth of the SB not only in Tr4 and Tr5 tracheal metameres, the two unique segments that develop during metamorphosis, but in all tracheal metameres (Fig. 1 A, D and E, F). Similar results were obtained using a constitutive activated form of the FGF receptor Btl (Torso-Btl) under the control of the same driver (Supplementary Fig. 1). The fact that only the Tr4 and Tr5 metameres develop during metamorphosis suggests that the FGF ligand Bnl might be expressed only in those metameres. To check this possibility, we analyzed the expression of Bnl in the larval trachea using a specific enhancer trap line that recapitulates its expression [16]. As expected, Bnl was only detected in the transverse connective branch of Tr4 and Tr5 at low levels at L2 (Fig 1G-G’) and higher expression during L3 (Fig. 1H-H’).

**Figure 1.**
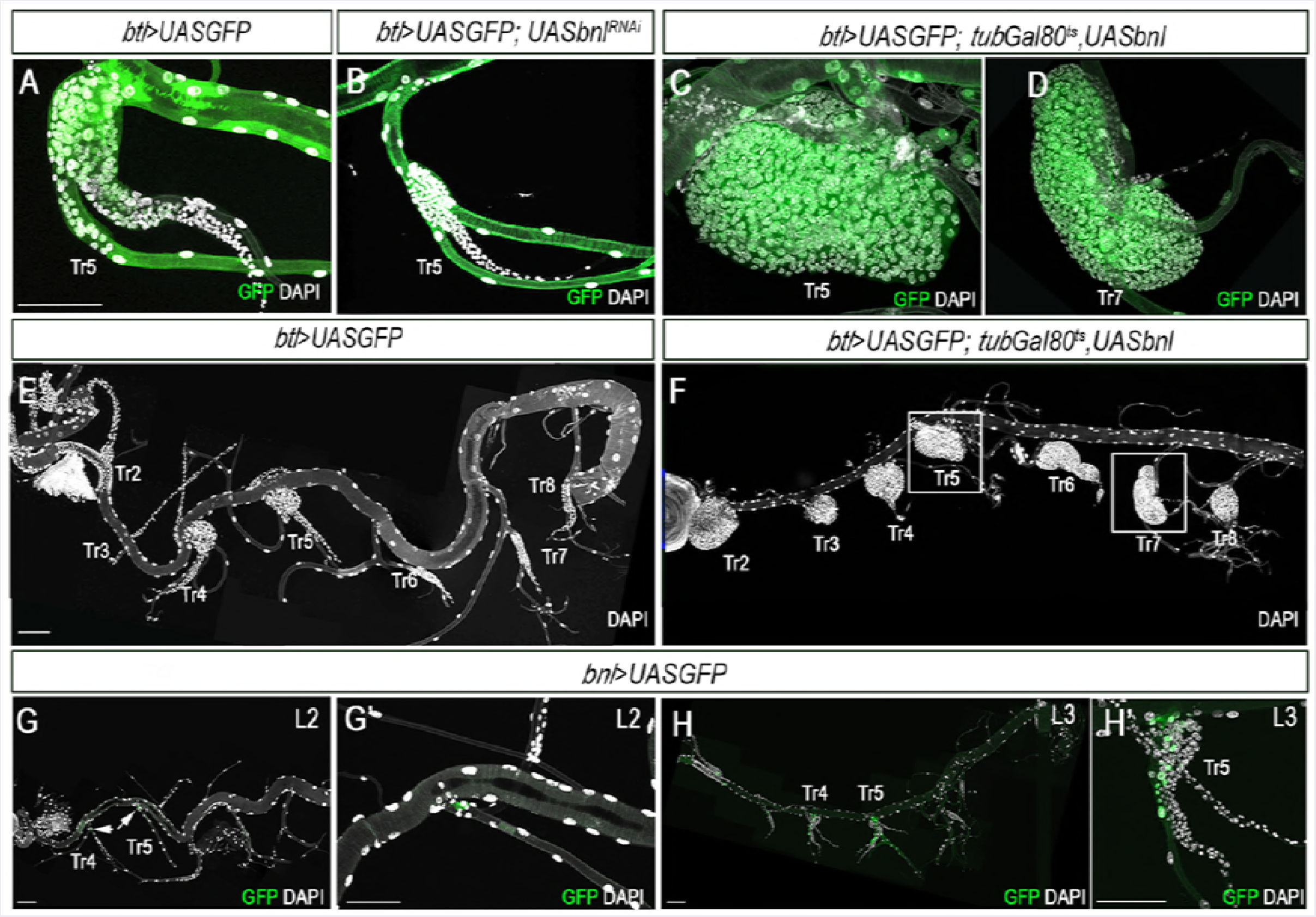
Bnl/FGF signaling activation initiates SB development. (A) *btl>UASGFP* control Tr5 SB, stained for GFP (green) and DAPI (grey). (B) Reduction in the number of tracheoblast cells in Tr5 SB of late L3 larvae upon depletion of FGF ligand Bnl. (C) T r5 and (D) T r7 SB cells overexpressing the FGF ligand *bnl* under the control of *btlGal4*. (E) *btl>UASGFP* control whole tracheal system showing all SB marked with DAPI. Note that only Tr4 and Tr5 SB develop. (F) Whole tracheal system of late L3 larvae overexpressing *bnl*. Note that over activation of Bnl/FGF signaling induces the development of all SB. (G-G’) Expression pattern of the *bnlGal4* reporter visualized by GFP (green) at L2 and (H-H’) late L3. DAPI is in grey. Scale bars represent 100 μm in all pictures.

We then analysed whether the activation of Bnl/Fgf signaling in the SB adult progenitor cells induces either cell growth or cell division. The highest rate of proliferation in the SBs takes place in the cells of zone 2 at mid-late L3 [13]. Using phospho-histone H3 (pH3), which labels mitotic cells in the G2/M transition [17], we measured the proliferation rate of zone 2 SB cells in mid and late L3 under different conditions of Bnl/Fgf signaling activity. In the control, PH3 positive cells were detected in zone 2 in mid and late L3, which later will generate a pool of differentiated cells in zone 1 (Fig. 2 A-A’’’ and G). In contrast, the inactivation Bnl/Fgf signalling by depletion of *bnl* under the control of *btlGAL4* resulted in a complete absence of mitotic cells and a reduced number of SB cells at late L3 (Fig. 2B-B’’’ and G). Consistently, the over-expression of the constitutive activated Torso-Btl receptor in the SB produced an increase of PH3 positive cells and the consequent dramatic increase of differentiated SB cells at late L3 (Fig. 2 C-C’’’ and G). We confirmed these results by clonal analysis. Thus, Btl dominant negative (Btl^DN^) overexpressing clones became visible in a very low frequency and small size than control clones (Fig. 2 D-E’ and H), whereas clone cells overexpressing Btl-Torso were bigger in size and cell number than wild type clones (Fig. 2 F-F’ and H). Altogether, these results strongly suggest that Fgf/Bnl signaling is necessary and required to induce the proliferation of adult progenitor cells in the SBs.

**Figure 2.**
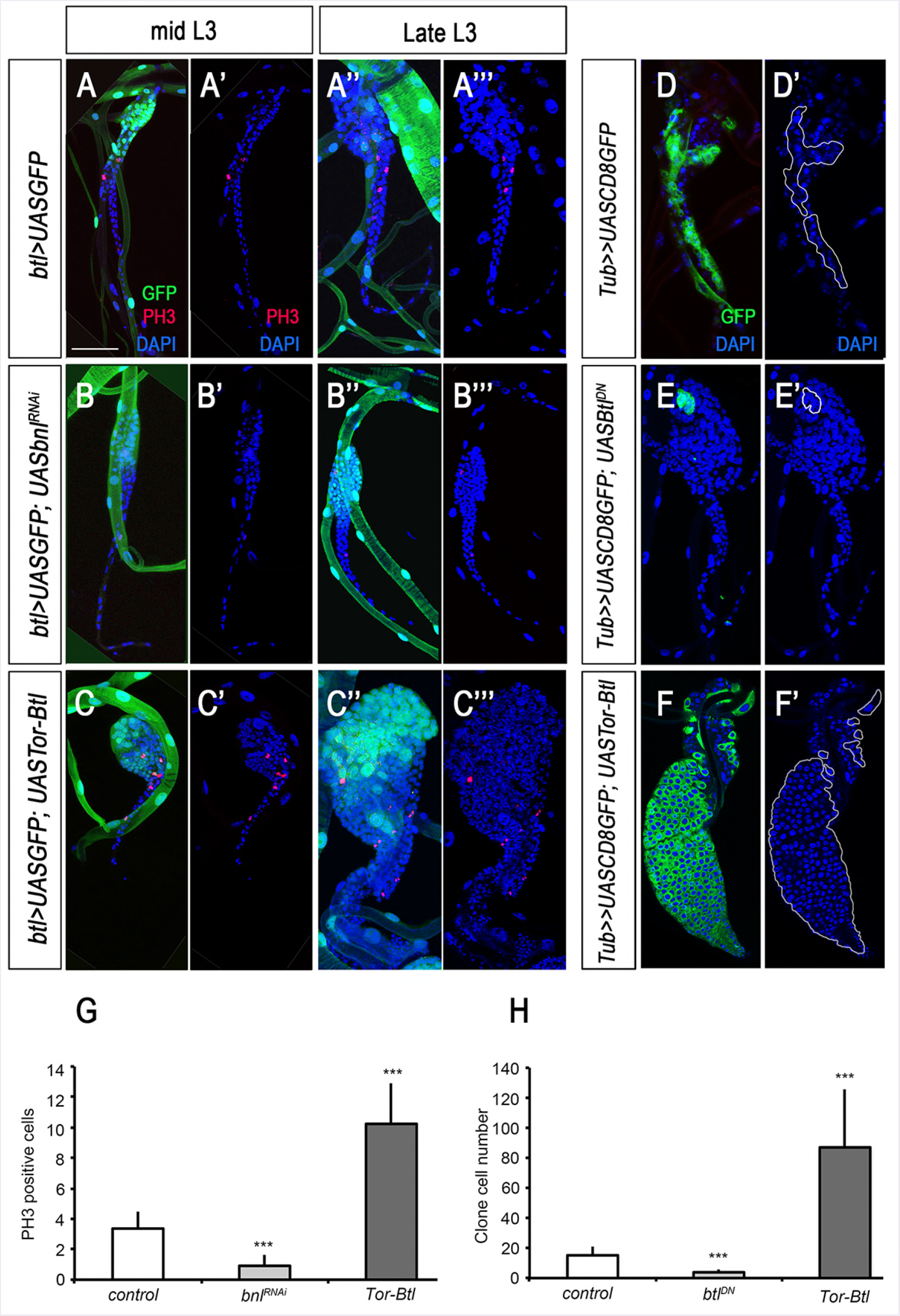
Bnl/FGF signaling induces SB adult progenitor cells proliferation. (A-A’’’) Control *btl>UASGFP* Tr5 SB of early and late L3 larva. (B-B’’’) Tr5 SB of early and late L3 larva depleted of *bnl*. Note the lack of PH3 positive cells compare to control. (C-C’’’) Tr5 SB of early and late L3 larva overexpressing a constitutive active form of the FGF receptor Btl (Tor-Btl*). (D-D’) SB with flip-out control clones visualized by GFP. (E-E’) SB with an overexpressing clone of a dominant negative form of the FGF receptor Btl (*UASBtl^DN^*). (F-F’) SB with an overexpressing clone of a constitutive active form of the FGF receptor Btl (Tor-Btl). In all pictures GFP is in green, PH3 in red and DAPI in blue. (G) Graph showing the average number of PH3-positive cells in the SBs of WT larvae and larvae ectopically expressing *UASbnl^RNAi^* or *UASTor-Btl\** under the control of *btlGAL4* (Student’ s t test, n>10 SBs; ***p < 0.0001). (H) Graph showing the average number of either *UASGFP, UASBtl^DN^* or *UASTor-Btl\** overexpressing clone cells (Student’ s t test, n>10 SBs; ***p < 0.0001). Scale bar represent 50 μm in A-C’’’ and D-F’.

Then, we investigated whether Bnl/Fgf signaling in tracheoblast proliferation requires transcriptional regulation. To address that, we analyzed the expression of *pnt*, the Ets domain transcription factor that mediates Fgf/Bnl signaling transcription activity in embryonic and larval tracheal cells [11]. Using a specific enhancer trap *pnt-lacz*, we found that *pnt* was specifically expressed in the cells of zone 2 and 3 of the SB, where Bnl/Fgf signaling presumably was active (Fig. 3 A-A’). To confirm that the expression of Pnt is related to Fgf/Bnl signaling activation, we overexpressed *pnt^RNAi^* in SB progenitor cells under the control of *CiGal4*, a specific driver of SB cells. Interestingly, we found that depletion of *pnt* impaired SB growth by reducing cell proliferation (Figure 3 B-C’). Similar results were obtained when overexpressing *pnt^RNAi^* flip-out clones were generated in the SB, as a low number of small clones were detected, suggesting that Pnt is required to mediate FGF signaling in the SB (data not shown). It is important to note, however, that Pnt also transduces the activation of the Epidermal Growth Factor (EGF) pathway, inducing mitotic division of the Air Sac Primourdium (ASP) tracheal cells through the phosphorylated isoform of Pnt, PntP2 [7,11]. To see whether this mechanism also operates in the SBs, *UAS-EGFR^RNAi^* was overexpressed under the control of the SB specific driver *CiGal4*. Interestingly, depletion of EGFR in the SB did not impair proliferation (Fig. 3 D-D’) suggesting that, in contrast to the ASP, SB cell proliferation is only controlled by the activation of the Fgf/Bnl signalling through Pnt.

**Figure 3.**
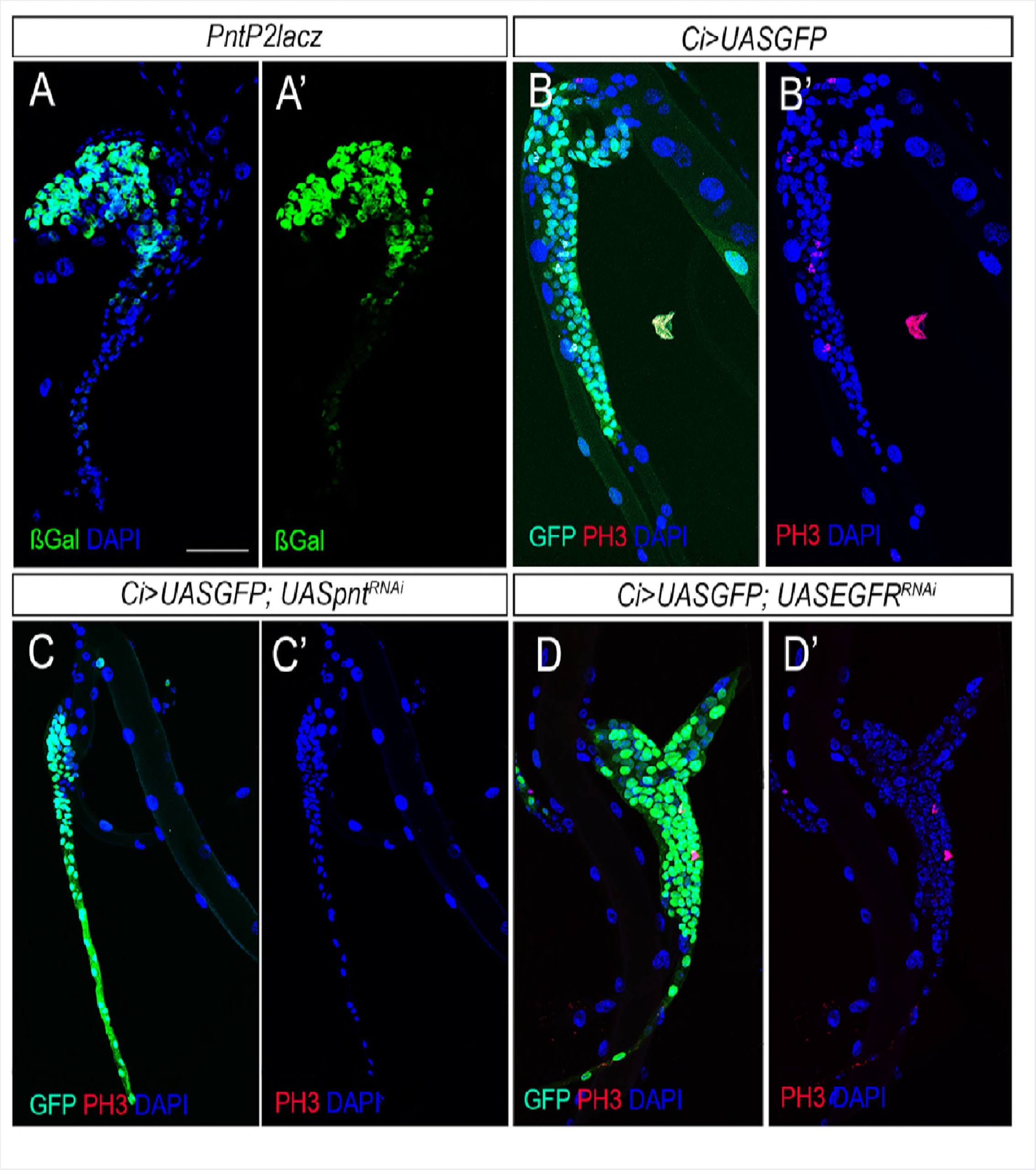
Bnl/FGF signaling acts on tracheoblast proliferation trough Pnt. (A-A’) Expression of the PntP2 reporter line in the SB visualized by anti β-Gal. (B-B’) Control *Ci>UASGFP* Tr5 SB. (C-C’) SB depleted of *pnt* by overexpression of *pnt^RNAi^* are unable to develop. (D-D’) Overexpression of *UASEgfR^RNAi^* under control of *CiGal4* in the SB. In all pictures GFP is in green, PH3 in red and DAPI in blue. Scale bars represent 50 μm in A-D.

### Bnl/Fgf signaling acts independently of the transcription factor *ct*

Our results above provide compelling evidence for the role of Bnl/Fgf signaling in promoting tracheoblast proliferation in the SBs. Previous studies, however, have proposed the transcription factor *ct* as the main factor that coordinates cell proliferation in the SB [13,15]. Therefore, one possibility is that Bnl/Fgf signaling may control proliferation by regulating *ct* expression. To solve this question, we over-activate or inactivate the Bnl/Fgf signaling pathway in adult progenitor cells of the SB and analysed the expression pattern of *ct* in those cells. Interestingly, under any of these conditions, *ct* expression was unaffected (Supplementary Fig 2), suggesting that Bnl/Fgf signaling promotes proliferation in Tr4 and Tr5 SBs without disturbing *ct* expression.

Then, we investigate whether Ct expression is necessary for Bnl/Fgf signaling activity to maintain proliferation of the SB cells. In fact, it has been shown that Ct restricts the expression of the FGF receptor Btl to cells in zone 1 and 2 of the SB [13,15]. According to this regulation, a reduction of Ct expression would induce an expansion of Btl expression into zone 3, promoting ectopic cell proliferation. Conversely, overexpression of Ct would repress Btl expression in zone 1 and 2, diminishing cell division. To verify this hypothesis, we either overexpressed or depleted Ct in the SB under the control of *btlGal4; tubGAL80^ts^* driver, rearing the animals at 25°C to avoid cell death as Ct acts as a cell survival factor in the SB [15]. As predicted, overexpression of Ct in SB cells reduced the total number of SB cells, probably due to reduction of Btl protein (Fig. 4 B-B’ and F). We confirmed this by rescuing the cell number defect caused by the overexpression of Ct by co-expressing the constitutive activated form of Btl, Torso-Btl (Fig. 4 CC’ and F). Similar effect was obtained when flip-out clones overexpressing *UAS-ct* and *UAS-Torso-Btl* were generated in the SB (Supplementary Fig. 3 A-B). In contrast, depletion of Ct in the SB cells increased total SB cell number (Fig. 4 D-D’ and F). If the positive effect of Ct knockdown in proliferation of the SB cells is due to a higher expression of Btl, then inactivation of the pathway would reduce the number of SB cells. As expected the inactivation of Btl pathway by coexpressing *UASbnl^RNAi^* in *ct* depleted tracheal cells reduced dramatically the number of SB cells (Fig. 4 E-F). Conversely, overactivation of the Bnl/Fgf pathway in SB clone cells depleted of *ct* resulted in a dramatic overproliferation (Supplementary Fig. 3 C-D), suggesting that the effect of Ct on proliferation depends on the regulation of Btl expression. Altogether, we conclude that Ct expression in SB cells is only required to restrict the population of SB cells that express Btl receptor, but not to initiate tracheoblast proliferation.

**Figure 4.**
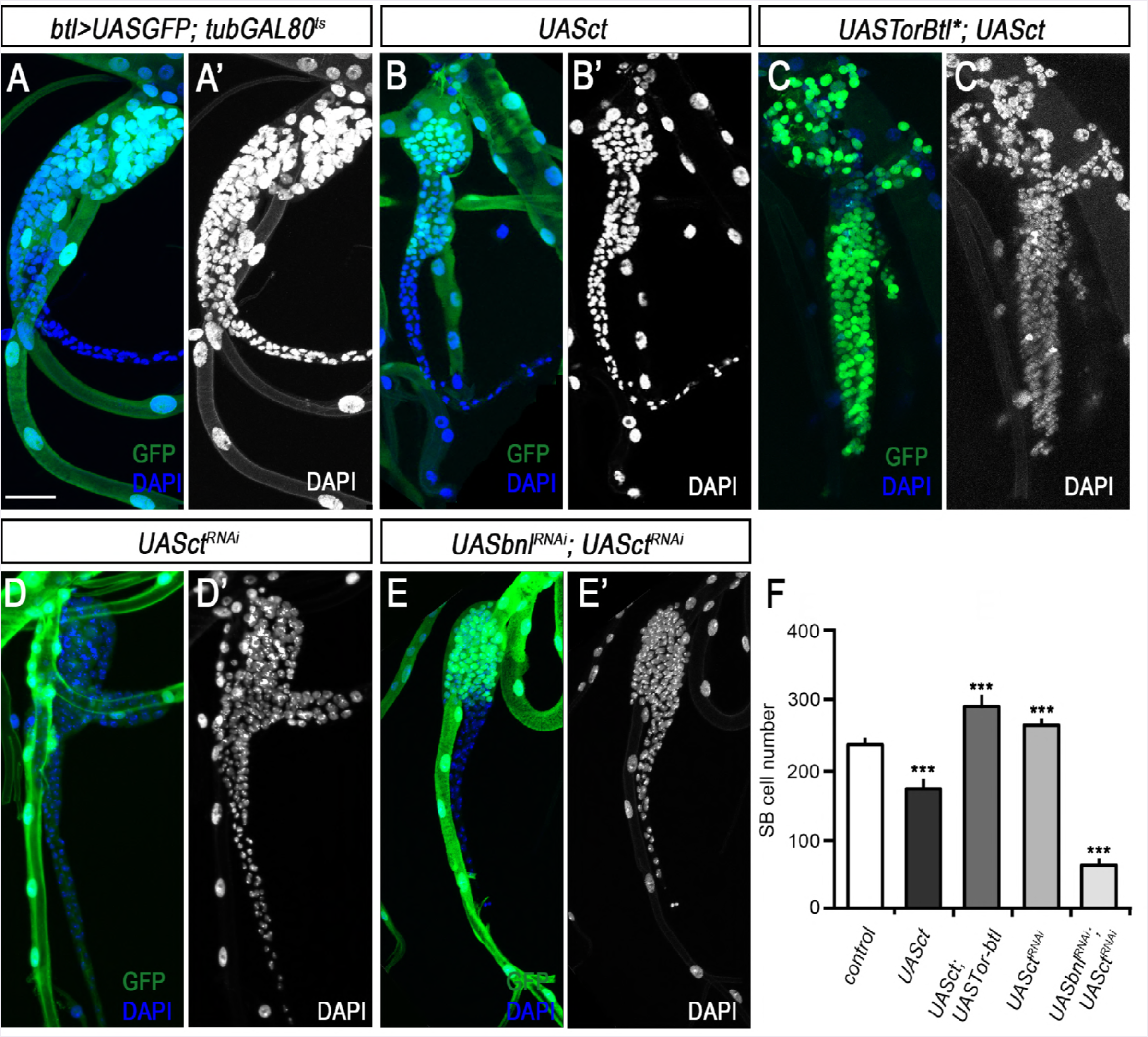
Bnl/Fgf signaling acts independently of transcription factor *ct*. (A) Control *btl>UASGFP; TubGal80^TS^* Tr5 SB of late L3 larva. (B-B’) Overexpression of Ct under control of *btl>UASGFP; TubGal80^TS^* in the SB. (C-C’) Rescue of induced Ct depletion proliferation defect by *UAS-Btl-Tor\**. (D-D) SB depleted of Ct by overexpression of *ct^RNAi^* (E-E’) Inactivation of Bnl/FGF signaling in SB cells depleted of *ct* are unable to proliferate. In all pictures GFP is green, and DAPI in blue and grey. (F-F’) Quantification of the total number of SB cells in the anteriorly described conditions compared to control (Student’ s t test, n>10 SBs; ***p < 0.0001). Scale bars represent 50 μm in A-E’.

### Bnl/Fgf signaling pathway promotes endoreplication in SB differentiated cells

As described above, activation of Bnl/Fgf signaling pathway in tracheoblasts of zone 2 induces cell proliferation. However, upon entering into zone 1, where Btl is expressed at high levels, these cells stop proliferation and initiate one round of endoreplication [13]. This transition depends on the repression of Ct by the Notch signaling, which allows the specific expression of the endocycle marker Fzr in SB cells of the zone 1 (Fig. 5A) [15]. Fzr is a Cdh1-like positive regulatory subunit of the APC/C that induces the degradation of the mitotic cyclins in G1 thus promoting endoreplication [18,19]. As Bnl/Fgf signaling pathway is active in zone 1 SB cells, it is possible that this pathway could also promote endoreplication. To understand the potential role of Bnl/Fgf signaling pathway in endoreplication of SB cells, we first analyzed the expression of *Fzr* in cells of zone 1 with overactivation and inactivation of the pathway. Inactivation of the pathway by depleting Bnl in all tracheal cells abolished the expression of *Fzr-lacZ* (Fig. 5 AB). However, we cannot discard that the impaired expression of Fzr was due to the fact that inactivation of Bnl/Fgf signaling in SB cells prevents the development of the SB. To avoid this problem, we generated flip-out clones overexpressing Btl^DN^, and found that clone cells with inactivated Bnl/Fgf signaling presented less expression of *Fzr-lacz* and smaller nucleus with less genomic DNA as measured by their chromatin value (C value), when compared to their control counterpart cells (Fig 5 C-C’’’ and I). In contrast, overactivation of Bnl/Fgf signaling by overexpression of Bnl in all tracheal cells increased the expression level of Fzr in SB cells that underwent one extra round of endocycle resulting in increase in the C value from 4C to 8C (Fig 5 D-D’ and J). A higher rate of endoreplication was also observed when flip-out clones overexpressing Btl-Torso were generated in the SB cells, as clone cells presented bigger nucleus and DNA content (8C) than the surrounding control cells (4C) (Fig 5 E-E’ and I). The results above suggest that the Bnl/Fgf signaling pathway promotes either proliferation or endoreplication of SB adult progenitor cells depending on the absence or presence of Fzr, respectively. If this were the case, depletion of Fzr would result in an increase of mitosis in zone 1 cells. Confirming this possibility, depletion of Fzr in the SB induced a switch from endoreplication to cell proliferation as detected by the detection of PH3 positive cells in zone 1 and the consequent reduction of cell size when *Fzr^RNAi^* was overexpressed under the control of *CiGAL4* (Fig. 5 F-G’). Consistently, over-activativation of Bnl/Fgf signaling activity in absence of Fzr dramatically increased the number of non-endoreplicative cells in zone 1 (Fig. 5 H-H’). Altogether, our results demonstrate that the Bnl/Fgf signaling pathway boosts cell proliferation in Fzr-absent SB cells of zone 2 and endoreplication in SB cells of zone 1 that express *fzr*.

**Figure 5.**
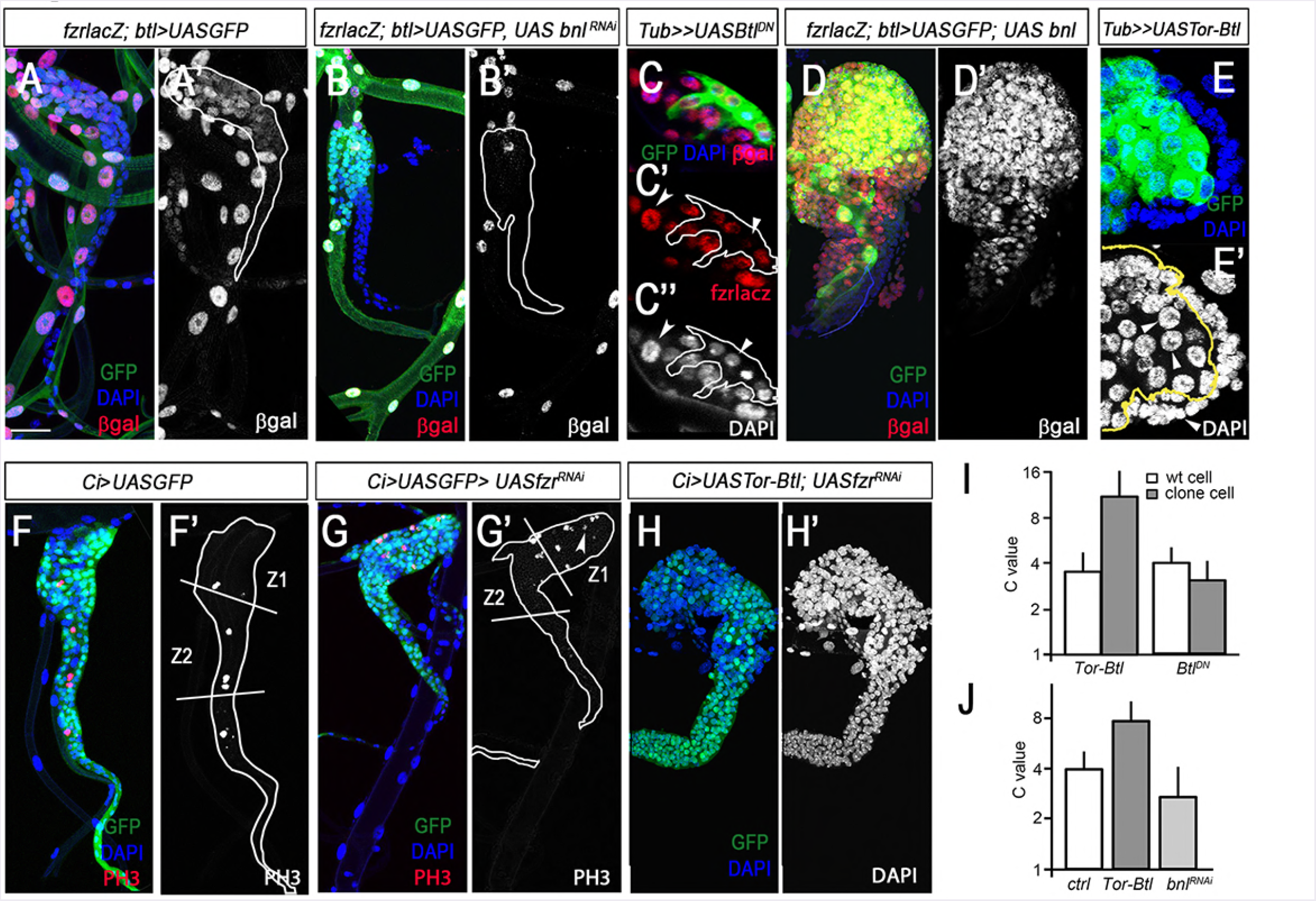
Bnl/Fgf signaling promotes endoreplication in SB differentiated cells. (A-A’) *fzr-lacZ* reporter expression in SB of late L3 larva (red) showing SB differentiated cells. (B-B’) *fzr-lacZ* expression in SB of late L3 tracheal system depleted of Bnl. (C-C’’) *fzr-lacZ* expression in clone cells overexpressing *UASbtl^DN^* visualized by GFP. (C’) Note the reduction of Fzr expression of clone cells and (C’‘) nuclear size compared to adjacent control cells (arrowheads). (D-D’) *fzr-lacZ* expression in SB cells overexpressing the FGF ligand Bnl. (E-E’) Flip-out clone overexpressing Tor-Btl marked by the expression of GFP. Increased of the nucleus size is observed in clone cells compared to the surrounding control cells (arrowheads). (F-F’) Control *CiGAL4 UASGFP* Tr5 SB of late L3 larva stained for mitotic marker PH3 (red and grey). (G-G’) SB cells depleted of *fzr*. Note the increase of PH3 positive cells in the SB differentiated cells of zone 1 (arrowhead). (H-H’) Overexpression of a constitutive active form of the FGF receptor Btl (Tor-Btl) in the Tr5 SB depleted of *fzr*. (I) The C value of clone cells in zone 1 of SB overexpressing either *Tor-Btl* and *Btl^DN^* compared to surrounding control cells. (J) Graph showing the C value of control zone 4 and zone 1 SB cells, and zone 1 SB cells overexpressing *Tor-Btl* and *bnl^RNAi^*. Scale bars represent 50 μm in A-B’ ‘, D-D’ and F-H’.

## Discussion

Our analysis of Bnl/Fgf signaling on SB morphogenesis shows that (i) the activation of the pathway in the Tr4 and Tr5 SBs is required and sufficient to initiate the normal development of these SBs, (ii) the action of Bnl/Fgf signaling promotes either proliferation or endoreplication of the SB cells depending on the expression of Fzr, and (iii) Ct regulates the SB cell behavior mode by targeting Fzr at the transcriptional level, but it is dispensable for Bnl/Fgf signaling activity. Our results illustrate how the activation of one signaling pathway, the Bnl/Fgf pathway, along the SB induces two different processes, cell proliferation and endoreplication, depending on the genetic context of the cell. Our data also demonstrate that the activation of Bnl/Fgf signaling, rather than the levels of Ct, controls the initiation of SB development and cell proliferation (Supplementary Fig. 4).

Bnl/Fgf signaling in *Drosophila* has been described as the main pathway that guide and differentiate tracheal cells during embryogenesis to form the larval trachea and also during metamorphosis to remodel and form the definitive adult trachea [2,3,5,7-13]. In addition to these well established roles, our results describe for the first time a role of Bnl/Fgf signaling in SB development by promoting cell proliferation and endoreplication. To date, the transcription factor Ct had been considered the factor that promotes either cell proliferation or differentiation in the SB depending on its abundance [15]. Our work, however, provides several lines of evidence demonstrating that it is not Ct but rather Bnl/Fgf signaling activity that is responsible for promoting SB cell proliferation and endoreplication: (1) SB cells proliferate even when Ct is depleted in these cells (Fig. 3 D); (2) absence of Bnl/Fgf signaling activity abolished SB development even in the presence of Ct expression; (3) ectopic activation of the Bnl/Fgf pathway induces SB growth by the dramatic increase of tracheoblast proliferation; (4) whereas Ct is similarly expressed in the SB of every tracheal metamere, Bnl is specifically expressed only in the Tr4 and Tr5 metameres, the only two metameres that will grow and develop (Fig. 1 F-G’); and (5) overexpression of *bnl* in all tracheal cells initiates the SB development of every SB (Fig 1 F). The factor that activates and restricts bnl expression in Tr4 and Tr5 is still unknown. The regulation of bnl expression is very complex as its expression in the ectoderm and tracheal cells is very dynamic during development [12,16,20]. Nevertheless, it is likely that Hox genes control the expression of *bnl* in the larva tracheal system. In this sense, Tr4 and Tr5 are specified by the expression of low levels of Abdominal A and Ultrabithorax [8,10,14], a hox code that may allow the expression of Bnl. Further experiments are needed to check this hypothesis.

Although Btl had been originally related to the proliferation of the ASP tracheoblasts [12], recent works have shown that Bnl/Fgf signaling promotes tracheoblast mitosis indirectly through the activation of the EGF ligand vein [7,11]. In contrast, our data shows that in the SB, Bnl/Fgf signaling promotes proliferation directly via the transcription factor *pnt*, and independently of the EGF pathway (Fig. 3). Therefore, our data show a direct role of Bnl/Fgf signalling in proliferation in *Drosophila*, in a similar way that occurs during the mammary gland development, where FGF signalling stimulates cell proliferation to generate cells both at the branching epithelium tips and cells in the subtending duct [21,22].

Contrary to previous reports, we also show that the main role of Ct in the SB is to determine the cell mode of tracheoblast by regulating the expression of Fzr. In this sense, our data indicate that Bnl/Fgf signaling induces cell proliferation or endoreplication depending on the presence or absence of Ct, respectively. Thus, Ct acts in the SB like in the ovary follicular epithelium where it also regulates the switch from mitotic cycles to endoreplication (Sun, 2005). The requirement for Ct in maintaining the mitotic cell cycle in *Drosophila* tracheoblast echoes its role in mammalian systems. The data in mammals suggest that CDP/Cut expression or activity might be restricted to proliferating cells [23]. Interestingly, the expression of the mouse CDP/Cut protein, Cux-1, in the kidney was found to be inversely related to the degree of cellular differentiation [24]. In addition, it has been shown that depletion of Cux-1 resulted in a significant increase in binucleate hepatocytes [25].

Our data indicates that Bnl/Fgf pathway not only initiates SB development and promotes cell proliferation of tracheoblast but also promotes cell endoreplication. In zone 1 of the SB, the activation of Notch signaling represses Ct expression thus allowing the initiation of endoreplication through the upregulation of the APC activator Fzr/Cdh1 [26]. Once activated, the successive endocycles are regulated by an intrinsic oscillator that consist of alternate APC activator and the levels of Cyc E [27]. Depending on the number of the times that the oscillator is on give rise to cells with 4C, 8C, 16C, 32C, etc. The activity of the Bnl/Fgf pathway seems to regulate the oscillator in SB differentiated cells, as inactivation of the pathway reduces the DNA content, whereas its overactivation increases the number of endocycles. Interestingly, this different effect of the signaling pathway promoting cell differentiation or endoreplication depending on the cell context is reminiscent of the EGFR signaling in the adult gut. After gut epithelial damage, EGFR signaling drives proliferation of intestinal stem cells (ISC), as well as endocycling in differentiated enterocits [28]. As EGFR and Bnl/Fgf pathway share many downstream components, including the transcription factor *pnt*, it is conceivable that the mechanism to promote endoreplication might be very similar. Another example is found in the oncogene *Dmyc*, which stimulates cell proliferation of ISCs in the Drosophila adult midgut [29] as well as endoreplication of fat body cells [30]. The hippo pathway has also been involved in promoting cell proliferation and endoreplication in larval tracheal cells depending on the expression of *fzr* [14]. However, the role of Bnl/Fgf pathway in SB development contrasts with the effect of the FGF4 in mammals. In this case FGF is only required to maintain trophoblast stem cells and therefore cell proliferation, as its inactivation drive the formation of Trophoblast giant cells that growth by endoreplication [31].

Our observations of the Drosophila tracheal system reveal that one signaling pathway can be used in a specific developmental process to induce both cell proliferation and endocycling, and that this capacity may be more common than has been generally appreciated not only in development but also in cancer. In certain contexts, cancerous cells use endoreplication as a path to drug resistance [32,33]. Interestingly, different evidences point to upregulation of the FGF/FGFR signaling as a mechanism of chemoresistance and radioresistance in cancer therapy [34]. Future studies should prove a possible link between FGF/FGFR system and endoreplication in tumors and promise insight into how to treat therapy resistance cancers.

## MATERIALS AND METHODS

### Fly stocks

Details for all strains genotypes can be obtained from flybase (http://flybase.org) or in references listed here. Conditional activation of either RNAi or gene expression was achieved using the *Gal4/Gal80^ts^* System [35]. To overexpress UAS transgenes either in all tracheal cells or in Spiracular branch cells *btlGal4UASGFP; tubGal80^ts^* or *CiGal4; tubGal80^ts^* was used respectively. Crosses were kept at 18 °C until late in L2 when larvae were shifted to 29 °C for 48 hours and dissected. The following stock flies where obtained from the Bloomington Stock Center: *btlGal4* (#8807), *tubGal80^ts^* (#7016), *UASpnt^RNAi^* HMS01452 (#35038), *UAScut^RNAi^* JF03304 (#29625), *UASEGFR^RNAi^* (#25781), *fzr-lacZ^G0326^* (#12241) and *pnt^1277^(pntP2-lacZ, #837). UAS-fzr^RNAi^* (#2550 and #25553), *UASbnl^RNAi^* (#101377 and #5730) and *UASpnt^RNAi^* (#105390) from VDRC and *CiGal4* [36], *UASbtl^DN^* [37], *bnlGal4* [38], *UAStor-btl* [39], *UASbnl* [40] and *UASct* [41] were given.

### Flip-out clones

Females of the genotype: *hsflp^70^; UASCD8GFP; tub>y^+^>Gal4* where crossed with the following UAS transgenic males: *UAStor-btl, UASbtl^DN^ and UASct* and were kept at 25°C until they reached early third instar stages. After a 30 min heat shock at 37°C, the larvae where transferred back to 25°C for 14-16h and dissected.

### Inmunohistochemistry

Larval trachea was dissected at third larval instar and fixed in 4% formaldehyde for 20 min. Incubations were performed o/n at 4°C with primary antibodies and during to 2 h at r.t. with secondary antibodies. Samples were mounted in Vectashield with DAPI (Vectorlabs). The following primary antibodies were used: anti-Ct (2B10, 1:100) and anti-βGalactosidase (401.a, 1:200) from Developmental Studies Hybridoma Bank, and anti-PH3 (1:500) from Cell Signalling Technology. Fluorescent conjugated secondary antibodies were obtained from Molecular Probes. Images were obtained with SP5 confocal microscope and processed with either Fiji or Photoshop CS4 (Adobe).

### DNA Quantification

For DNA quantification, DNA staining intensity in the SB cells was obtained from z-stacked images of DAPI stained tracheal system. DNA staining intensity of SB cells of zone 1 was normalized using average DNA staining intensity in the SB zone 4 cells: DNA staining intensity in the SB cells of zone 1/DNA staining intensity in SB cells of zone 4. The C value of the control SB cells of zone 1 cells at late L3 was set to 4C [42].

## Acknowledgements

We thank Bloomington and VDRC for fly stocks and colleagues in the lab for discussions and reading of the manuscrit. Support for this research was provided by the Spanish MINECO (grant BFU2009-08748 CGL2014-55786-P to D.M. and X.F-M.) and by the Catalan Government (2014 SGR 619 to D.M. and X.F-M.). The research has also benefited from FEDER funds. J. Cruz was supported by CGL2014-55786-P from the Spanish MICINN, and C.d.M. was supported by FPI from the Spanish MICINN.

---

**Supplementary Figure 1.** Effect on SB development upon Bnl/FGF signaling overactivation (A) *btl>UASGFP* control whole tracheal system showing all SB marked with GFP (green) and DAPI (blue). Note that only Tr4 and Tr5 SB develop. (B) Whole tracheal system of late L3 larvae overexpressing the constitutive active form of Btl *(Tor-Btl*)*. Note that over activation of Bnl/FGF signaling induces proliferation of all SB. (C) Tr5 and (D) Tr7 *btl>UASGFP* SB control cells. (E) Tr5 and (F) Tr7 SB cells overexpressing the *Tor-Btl\** under the control of *btlGal4*. Scale bars represent 100 μm.

**Supplementary Figure 2. Bnl/FGF signaling effect on Ct expression** In all pictures GFP is shown in green, Ct in red and DAPI in grey (A-A’) *btl>UASGFP* control SB. (B-B’) SB cells overexpressing *Tor-Btl\**. (C-C’) SB cells overexpressing *bnl^RNAi^*. Scale bars represent 50 μm.

**Supplementary Figure S3. Effect of Ct in Bnl/FGF signaling activation** (A) Flip-out clones overexpressing *ct* marked with GFP and stained for Ct (red) and DAPI (blue and grey). (B) Flip-out clones co-overexpressing *UAS-Tor-Btl \** and *ct*. Note the bigger size of the clones. (C) Flip-out clones marked with GFP and DAPI (blue and grey) overexpressing either *ct*^RNAi^or (D) *Tor-Btl*\*and *ct^RNAi^*. Scale bars represent 50 μm.

**Supplementary Figure S4. Model depicting the spatial activation of FGF signaling in adult progenitor cells**. FGF signaling is activated by the ligand Bnl specifically in Tr4 and Tr5 tracheal metamers. The different outcome of the activation of FGF pathway, depending on the presence or absence of Ct allowing proliferation and endoreplication respectively, is also shown. In metameres Tr6 to Tr9 despite the presence of Ct, FGF signalling is not activated and therefore adult progenitor cells do not proliferate.

